# A novel nanobody-based approach for targeting heterogeneous *Acinetobacter baumannii* isolates and closely related pathogenic *Acinetobacter* spp

**DOI:** 10.64898/2026.05.06.723352

**Authors:** Anke Breine, Elena Jooris, Adam Valcek, Steven Van Meerbeek, Els Pardon, Delphi Van Haver, Evy Timmerman, Francis Impens, Jan Steyaert, Han Remaut, Inge Van Molle, Mihaela Gheorghiu, Daniela Tudor, Sorin David, Eugen Gheorghiu, Charles Van der Henst

## Abstract

*Acinetobacter baumannii* is a top-priority, ESKAPE pathogen that poses a major challenge to human health. The pathogen is difficult to combat due to its extensive arsenal of antibiotic resistance and its protective polysaccharide capsule. In addition, *A. baumannii* isolates are highly heterogeneous, which complicates the development of rapid detection methods or novel targeted therapeutic approaches. Here, we discovered and characterized a new biotechnological tool, the nanobody H7 (NbH7), along with its conserved target, the surface-exposed Omp25 protein of *A. baumannii*, and elucidated their interaction at the molecular level. Moreover, we demonstrate that NbH7-functionalized magnetic beads enable selective and efficient capture of *A. baumannii* from bacterial mixtures, including non-pathogenic intestinal bacteria. This provides proof of concept for a new targeting system that remains effective across diverse *A. baumannii* clinical isolates and capsule types and holds potential for use in diagnostic cell enrichment and targeted therapies.

## INTRODUCTION

Antimicrobial resistance (AMR) is a major global health threat. Due to its impact on mortality and morbidity rates and the associated healthcare costs, the World Health Organization (WHO) and the Centers for Disease Control and Prevention (CDC) have each compiled lists of top-priority pathogens for which therapeutic alternatives and more effective treatments are urgently required.^12^ The top priority pathogens share a substantial overlap with the ‘ESKAPE’ group of pathogens (*Enterococcus faecium, Staphylococcus aureus, Klebsiella pneumoniae, Acinetobacter baumannii, Pseudomonas aeruginosa*, and *Enterobacter* spp.), a group infamously known for representing major therapeutic challenges.^3^

Carbapenem-resistant *A. baumannii* (CRABs) is part of the ESKAPE group and the critical priority pathogen lists. *A. baumannii* is a human pathogen that causes infections at various anatomical sites and thrives in clinical settings.^4^ The pathogen has acquired its priority position due to a dangerous combination of characteristics, such as: (i) the facile acquisition of drug resistance genes, and (ii) its ability to survive for extended periods of desiccation.*A. calcoaceticus* and *A. pittii* are also recognized as pathogenic or newly emerging pathogenic species of *Acinetobacter* spp., so targeting these species is also relevant.

In recent years, traditional efforts to fight CRABs have failed^1^, emphasizing the need for alternative treatments. Any therapy for AMR should not only be effective but also reduce the selective pressure on the bacteria to develop resistance. The approach to counter AMR must be twofold: the development should focus on pathogen-specific therapeutics to reduce secondary effects, and on accurate, rapid diagnostic techniques to enable correct, early treatment decisions and prevent improper antibiotic use. Developing targeted therapies against *A. baumannii*, however, is complicated by the bacterium’s highly heterogeneous nature, driven by a small core genome and considerable genomic variability among strains.^5^ Identifying conserved molecular targets for this pathogen, along with the physico-chemical barrier generated by the protective polysaccharide capsule, has been a major challenge in developing targeted interventions and bacterial cell enrichment for early diagnostics. It is therefore essential to find novel targeted approaches and alternative strategies.

The use of Nanobodies (Nbs) is a valuable alternative approach for developing targeted therapies and rapid diagnostic techniques. Nbs, also known as VHH or single-domain antibodies, are the variable domain of a heavy-chain-only antibody found in camelids.^6^ These small proteins are remarkably stable, retain a high affinity and specificity for the antigen, and their modularity allows for the creation of multivalent constructs and fusion with other proteins.^7^ In the realm of targeted therapeutics and diagnostic tools, these are valuable characteristics. The implementation of Nbs has been established in numerous settings, including several studies on *A. baumannii*.^8,9^ However, whilst generating an Nb binder is possible, finding an Nb capable of binding diverse *A. baumannii* strains while retaining species specificity has proven to be the bottleneck.^10,11^

In this work, we identify a novel Nb targeting a conserved outer membrane protein of *A. baumannii*, characterize its molecular interactions, and demonstrate its promising early diagnostic potential through immunomagnetic capture of *A. baumannii* from a complex solution.

## RESULTS

### Identification of *A. baumannii*-specific nanobody NbH7

To explore targets on the cell surface of *A. baumannii* in an unbiased way, a nanobody (Nb) library was created by immunizing a llama with intact, inactivated *A. baumannii* bacteria. The immunization was done with the AB5075-VUB strain (referred to as AB5075C+), a clonal isolate of the broadly used AB5075 strain. ^12^ This strain is known to be virulent, capsulated and multidrug-resistant, including carbapenem-resistant, making it a relevant research strain.^13,14^ In addition, its non-capsulated mutant derivative (referred to as AB5075C-) was also used.^12^

The enriched Nb library was screened for binders recognizing diverse *A. baumannii* isolates with varying capsule types and densities: three isolates from our collection (AB3-VUB, AB180-VUB and AB220-VUB) and a clonal isolate of the frequently used ATCC17978^15^ (ATCC17978-VUB) strain.^16,15^ Binding was assessed on live cells using fluorescence microscopy with fluorescently labelled Nbs (see material and methods). A control Nb (CtrlNb) was included to account for aspecific interactions (**Figure S1**). Among the candidate binders, NbH7 showed consistent binding to all tested *A. baumannii* strains, including the highly capsulated, pandrug-resistant (PDR) strain AB3-VUB (**Figure 1A**).

**Figure 1.**
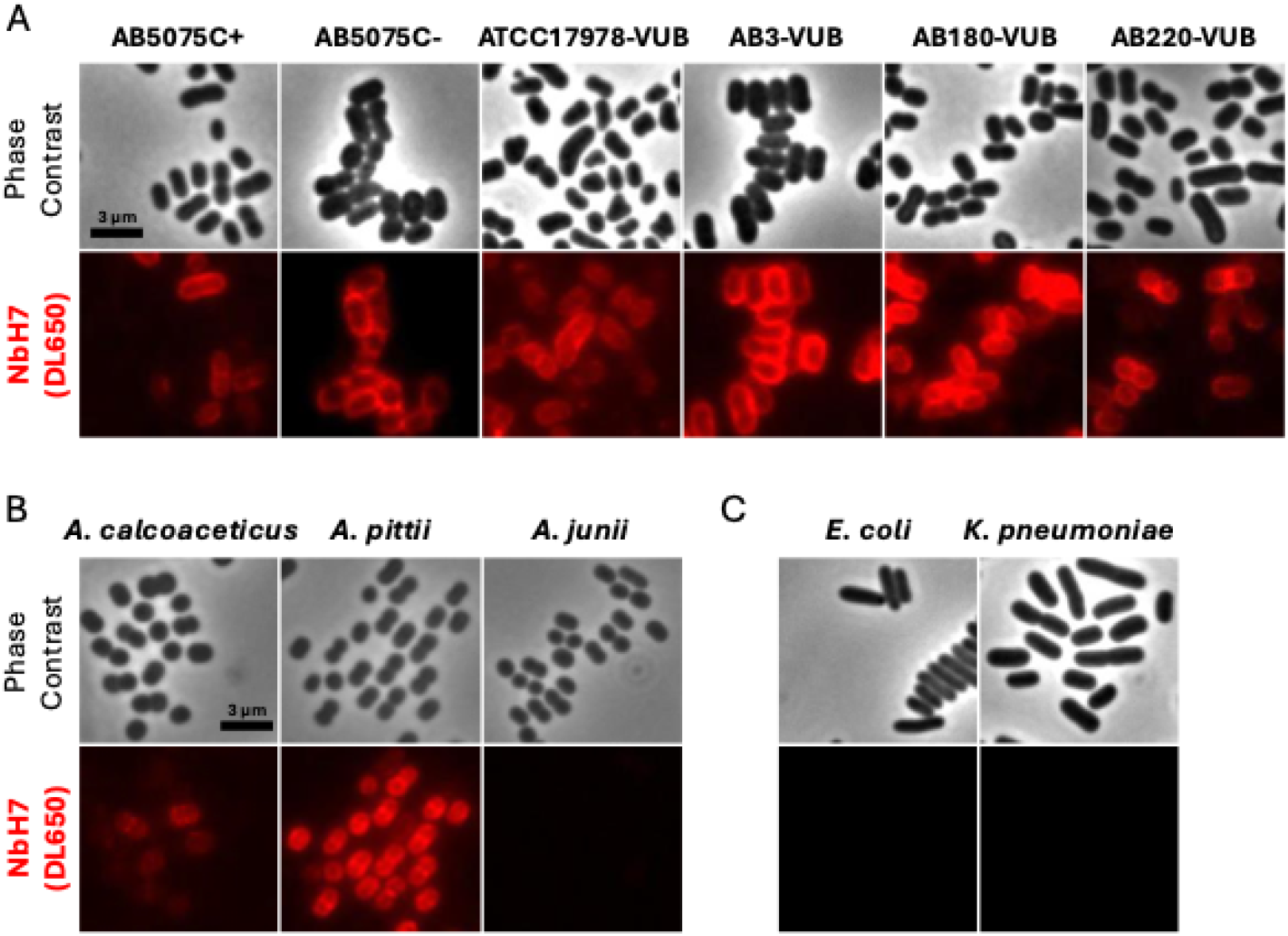
NbH7 is specific to a range of *Acinetobacter* strains. **A**. NbH7 was fluorescently labelled with DL650 and its binding was observed with fluorescence microscopy on different bacterial strains: six *A. baumannii* strains, of which “AB5075C+” is the WT capsulated strain and “AB5075C-” is the *itrA* non-capsulated deletant strain. Three other *Acinetobacter* species were also tested (**B**), as well as *Escherichia coli* S17 and *Klebsiella pneumoniae* (**C**). Scale bars: 3 μm.

NbH7 furthermore displayed a uniform labeling pattern on *A. baumannii*. Fluorescence intensity profiling of NbH7 on an *A. baumannii* strain constitutively expressing GFP in the cytoplasm revealed a membrane labelling pattern of bound NbH7, while the intensity profile of GFP shows cytoplasmic distribution (**Figure S2**). These results indicate that NbH7 binds to the cell surface of *A. baumannii*. Moreover, the homogeneous distribution and clear signal indicate that the Nb target is distributed across the entire cell surface rather than localized in discrete regions.

The binding range of NbH7 was further explored by testing its ability to bind to closely related species, such as *Acinetobacter calcoaceticus, Acinetobacter pittii* and *Acinetobacter junii*, but also less closely related species such as *Klebsiella pneumoniae* and *E. coli*. Fluorescence micrographs revealed that NbH7 binds to two out of the three tested *Acinetobacter* spp. (**Figure 1B**). For the tested *E. coli* and *K. pneumoniae* strains, no binding is observed (**Figure 1C**).

### NbH7 targets the conserved outer membrane protein Omp25

The target of NbH7 was identified by means of a pull-down assay performed on *A. baumannii* cell lysates. A protein band of approximately 25 kDa appeared in the NbH7 sample compared to the control sample (**Figure 2A**). Mass spectrometry analysis identified this protein as Omp25 (**Table S1**). To verify whether Omp25 is the target of NbH7, an *omp25* gene knockout of the AB5075 strain was generated. Binding analysis revealed a complete loss of NbH7 binding to this mutant strain (**Figure 2B**). Remarkably, recombinant expression of *A. baumannii* Omp25 in *E coli* results in NbH7 binding (AbOmp25) in *E. coli* (**Figure 2C**). This heterologous complementation further confirms that NbH7 specifically targets Omp25.

**Figure 2.**
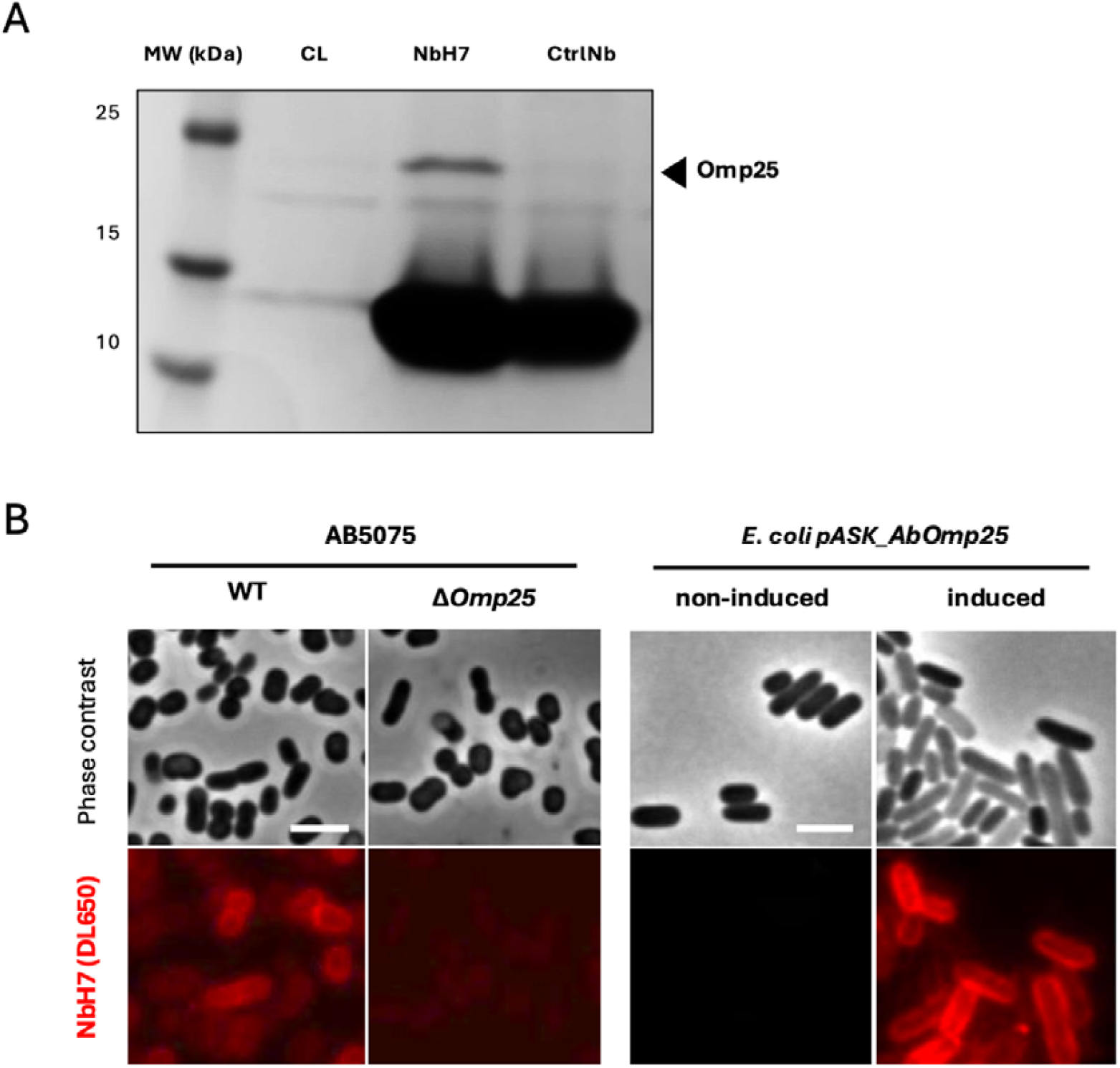
NbH7 binds the conserved outer membrane protein Omp25. **A** SDS-PAGE analysis and Coomassie blue staining of the pull-down using NbH7 and a control Nb (CtrlNb) on the cell lysate (CL) of *A. baumannii* (AB5075C-). CtrlNb is an unrelated Nb. The arrow indicates the additional band compared to the control. **B** Fluorescence micrographs of labelled NbH7 on *A. baumannii* and its Δ*omp25* derivative (left panel). Fluorescence micrographs of labelled NbH7 binding on *E. coli* recombinantly expressing (induced) or not expressing (non-induced) AbOmp25 (right panel). The scale bar indicates 3 μm.

Genomic conservation analysis indicates that the *omp25* gene is present in 1034 of 1035 (99.9%) complete *A. baumannii* genomes. The nucleotide identity ranged from 98.31% to 100% and the genetic environment of the *omp25* was conserved (**Figure S3**). Additionally, the comparison of their protein sequences showed that the different *A. baumannii* strains differ only in the signal peptide of Omp25. The *omp25* gene as well as the corresponding mature protein is thus highly conserved (**Figure S4**).

The genomic sequences of the three *A. calcoaceticus, A. pittii*, and *A. junii* strains tested are unavailable; thus, we could not directly compare Omp25 protein conservation levels across these related species. Nonetheless, to evaluate Omp25 conservation in these 3 species, the AbOmp25 amino acid sequence was blasted against the incomplete proteome of *A. calcoaceticus, A. pittii* and *A. junii*. One representative strain was chosen for each occurring percentage sequence identity for protein sequences of the same length (**Table S2**). Interestingly, *A. junii* has a homologous Omp25 sequence, even though no binding of NbH7 was observed on this strain. An amino acid sequence identity of approximately 93-96% for *A. calcoaceticus*, 94-96% for *A. pittii* and 81% for *A. junii* was observed. The lower conservation of the Omp25 amino acid sequence in *A. junii* can explain the lack of binding of NbH7. A multiple sequence alignment of the Omp25 protein sequences from these strains revealed that most mutations are clustered in the same regions (**Figure S5**).

### NbH7 targets the surface-exposed region of Omp25

To characterize the molecular interaction of NbH7 with Omp25, the protein complex was co-crystallized. The crystals of Omp25 in complex with NbH7 diffracted to 2.91 Å and could be used to obtain the experimental structure of the complex.NbH7 binds to the extracellular side of Omp25 (**Figure 3A**) with an interaction surface of 1010.4 Å^2^. The CDR3 of the Nb protrudes into the barrel and establishes its main interaction through residues GLU113, SER102 and TYR105, which form hydrogen bonds with corresponding residues of Omp25 inside the barrel **(Figure 3B)**. CDR2 and CDR1 both interact with loop residues **(Figure 3C, 3D)**, further stabilizing the interaction.

**Figure 3.**
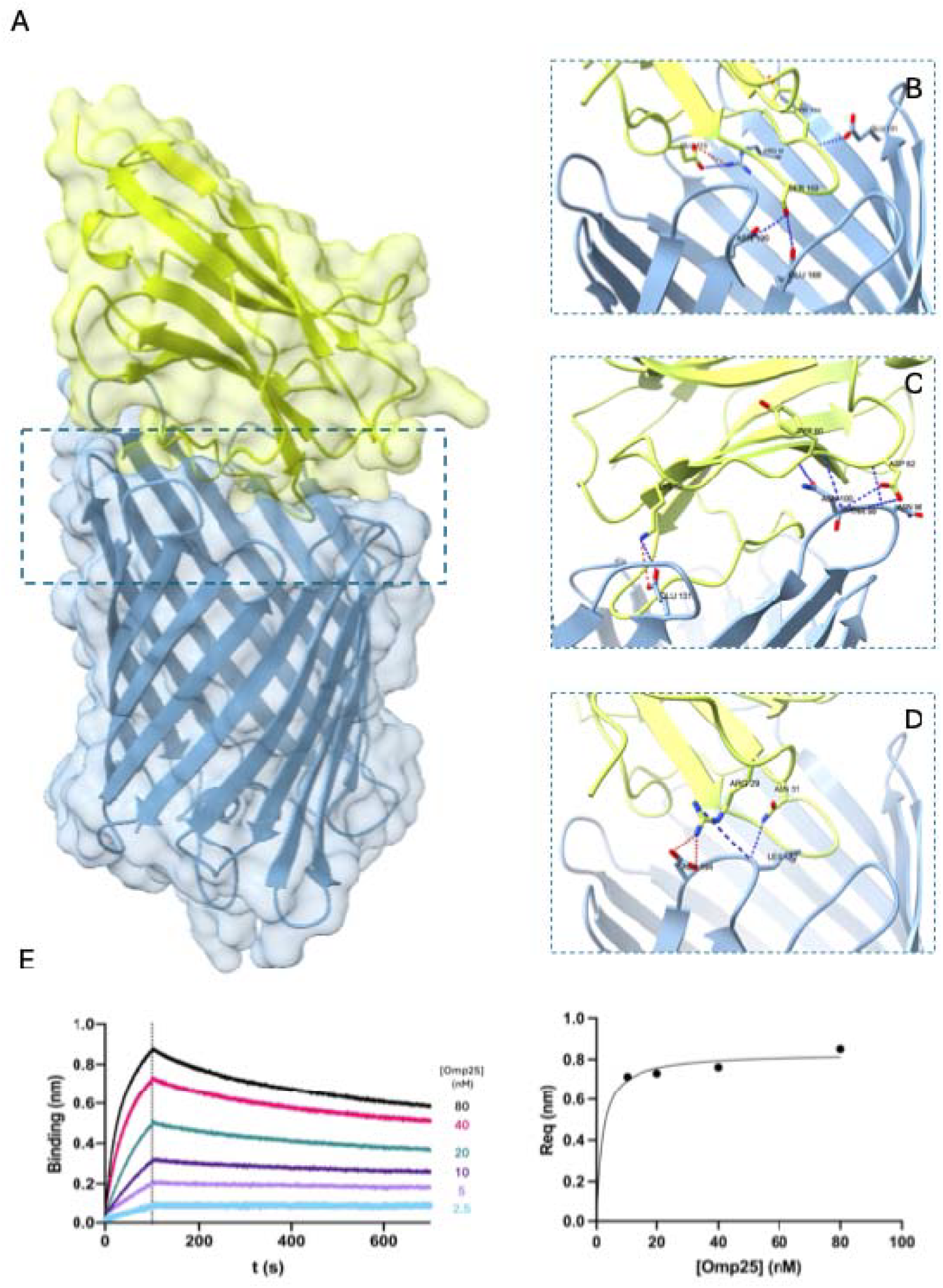
**A** Ribbon presentation of Omp25 in complex with NbH7. The structure was solved by X-ray crystallography at 2.91Å. Omp25 is colored in blue, NbH7 in light green. Interacting residues are indicated for the Nb CDR3 (**B**) CDR2 (**C**) and CDR1(**D**), hydrogen bonds are indicated in blue, and salt bridges in red. NbH7 NanoSaurus^18^ entry [SD-CVAP] **E** *Left panel:* Association and dissociation curves for the Nb with a two-fold dilution series of Omp25 concentrations. *Right panel:* The R_eq_-values plotted against the Omp25 concentration and fitted on a one-site specific binding curve to determine the equilibrium dissociation constant (K_D_ ± standard error). The R_eq_-values were calculated using Octet® Analysis Studio software and the graphs were plotted using Graphpad Prism.

To better understand this molecular interaction, NbH7 and Omp25 were recombinantly produced and purified (**Figure S6**). Their interaction was biochemically characterized using bio-layer interferometry (BLI), with NbH7 was loaded onto a Ni-NTA sensor and tested for binding against different Omp25 concentrations. This revealed a dissociation constant in the low nanomolar range (K_D_ of 7.69±0.74 nM), consistent with high-affinity binding (**Figure 3E**).

### NbH7-assisted immunomagnetic capture of *A. baumannii*

The application potential of NbH7 was explored by assessing its ability to capture *A. baumannii* from homogeneous and mixed bacterial samples when coupled to magnetic beads. Upon binding of target bacteria, these magnetic beads form clusters that can be monitored qualitatively (optically) and quantitatively (electrically), enabling rapid assessment of capture efficiency, specificity, and bacterial viability.^19^

Large bead clusters were observed **(Figure 4A)** when NbH7-coated magnetic particles (MPs) were incubated with the target bacterium *A. baumannii*. In contrast, only small or no clusters formed in the presence of the non-target bacterium *E. coli* **(Figure 4B)**, or in the absence of bacteria **(Figure 4C**), confirming NbH7-specific aggregation (**Figure S7**). In mixed samples of *A. baumannii* and *E. coli*, large clusters were still observed **(Figure 4D)**, indicating that the target bacterium *A. baumannii* dominates cluster formation even in mixed populations.

**Figure 4.**
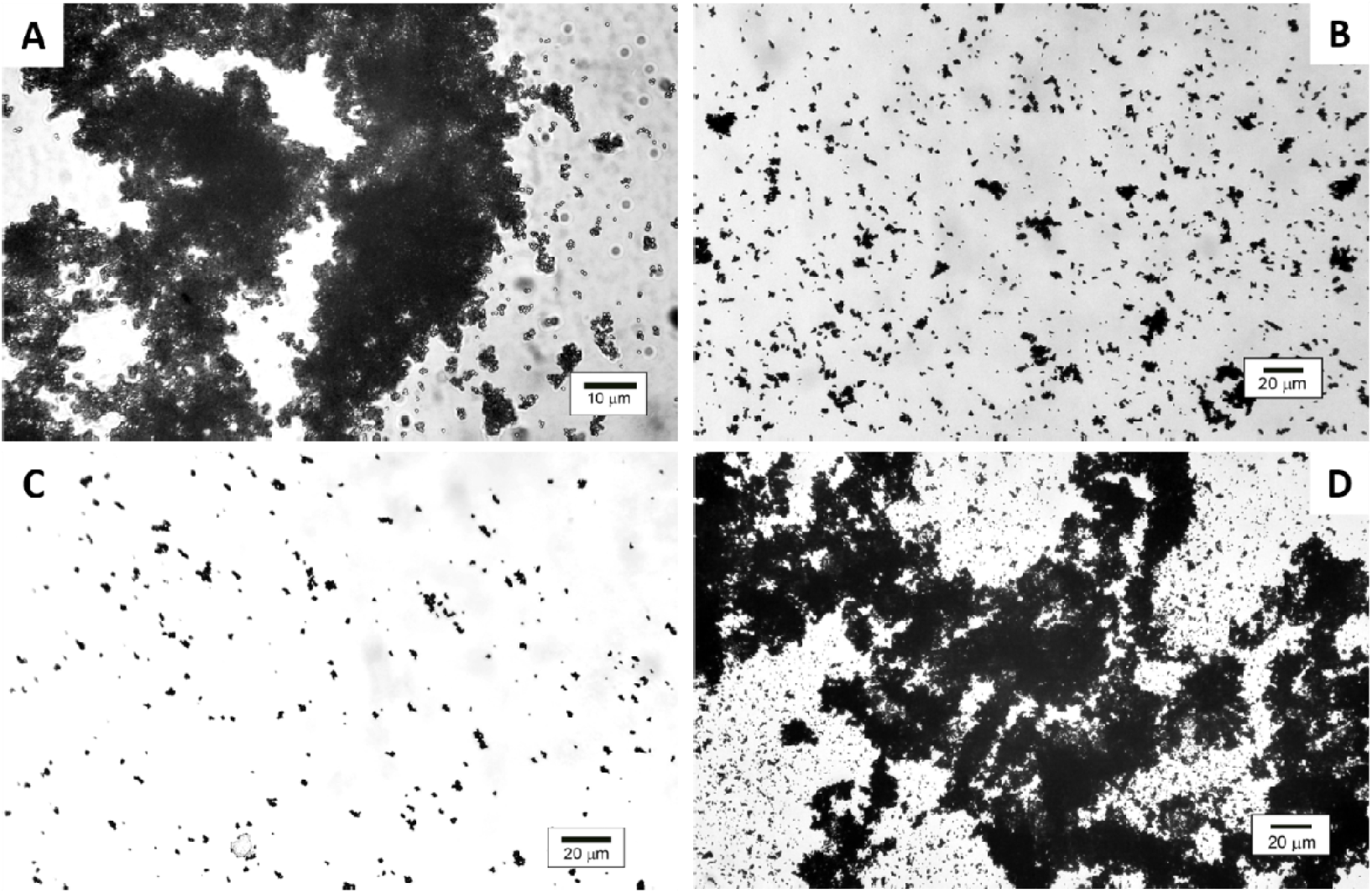
Cluster formation analysis by optical microscopy. (**A**) Transmission microscopy image of NbH7-coated MPs incubated with *A. baumannii*, showing the specific clusters and high aggregation potential. (**B**) Transmission images with non-target bacteria (*E. coli*), where very small, isolated clusters are observed, compared to (A). (**C**) Control transmission image of MPs in the absence of cells, similar to that obtained with non-target bacteria. (**D**) Characteristic transmission image of NbH7-coated MPs clustering in a mixed suspension of *A. baumannii* and *E. coli*.

Electrical analyses further quantified the cluster observations. A concentration-dependent increase in the normalized parameter Z_norm_ for *A. baumannii* compared to unbound MPs was revealed (**Figure S8A**). Significant increases in Z_norm_ were observed at 5 x 10^5^ and 10^6^ cells/mL, whereas lower concentrations did not differ significantly from controls (p < 0.05). No significant change was detected for *E. coli*, which produced signals comparable to MPs alone (**Figure S8A**).

The specificity of the cluster formation was tested under competitive conditions by mixing increasing concentrations of *A. baumannii* (10^5^ – 10^6^ cells/mL) and a fixed, excess amount of *E. coli* (10^6^ cells/mL).^20^ A concentration-dependent increase in the normalized parameter Z_norm_ was observed for *A. baumannii* compared to non-specific MPs under these conditions as well (**Figure S8B**). In addition, non-specific clustering was observed in all samples as well, including the control samples. This is a relevant indicator of the specificity and selectivity of the NbH7 affinity coating. **Figure S8B** shows that the NbH7-functionalized MPs selectively recognize and bind the target bacterium *A. baumannii* also in mixed cultures.

## DISCUSSION

*Acinetobacter baumannii* is infamous for its extensive genome plasticity and capsule diversity. This presents a central challenge in targeting conserved and surface-accessible antigens on this priority pathogen. In this study, we addressed this challenge by generating an unbiased nanobody library targeting the entire cell surface of a relevant *A. baumannii* isolate, without prior bias toward specific antigens. We selected nanobodies binding a highly diverse set of isolates representing this pathogen with a small core genome (approximately 16.5 %)^5^ and high variability. We identified NbH7 as a binder retaining its binding ability in these diverse isolates regardless of the capsule density, thickness, or type and the binding ability of the Nb.

The binding profile of NbH7 indicated that its target is homogeneously distributed across the entire cell surface rather than localized in discrete regions. This widespread distribution is advantageous for both diagnostic applications and targeted therapeutic strategies. Additional binding assays showed that NbH7 does not bind *A. junii, E. coli*, or *K. pneumoniae*. The lack of binding to *A. junii* might be a consequence of the absence of the target of NbH7 in *A. junii*, the presence of an altered target, or the presence of factors impeding the binding of NbH7. Nevertheless, this binding assay demonstrated the specificity of NbH7 for *A. baumannii* and closely related *Acinetobacter* species. The fact that NbH7 can recognize *A. baumannii* as well as *A. calcoaceticus* and *A. pittii*, the latter species recognized as pathogenic or emerging pathogens, is relevant. However, a discriminatory step in the diagnostic effort would be necessary to precisely identify which *Acinetobacter* spp. a patient is colonized or infected with.

The identification of the NbH7 target led us to discover an accessible, surface-exposed antigen: the outer membrane protein Omp25. Genetic deletion of *omp25* abolishes binding, while heterologous expression in *E. coli* restores it, demonstrating that Omp25 is both necessary and sufficient for recognition within the tested conditions. Remarkably, the *omp25* gene is part of the core genome, and the corresponding mature Omp25 protein sequences of the selected strains are identical. Structural characterization of the Omp25-NbH7 complex further detailed this interaction at the molecular level and provided insight into the binding interface.

Whether NbH7 has an intrinsic effect on *A. baumannii* remains to be elucidated. The structure of the complex confirms that Omp25 adopts a β-barrel conformation, as was proposed but not yet structurally confirmed by Siroy et al.^17^ The Nb binds to the relatively small protein (25 kDa) with its CDR3 protruding into the barrel. This binding mode suggests the Nb’s potential to modulate channel function, although no direct functional effects were assessed in this study. Previous studies have linked Omp25 and related outer membrane proteins to membrane permeability and stress responses, but their precise physiological roles remain unresolved.^17, 21,22^ Omp25 has a conserved motif, _43_PLAEAAFL_50_, that encodes an ⍰-helix and is present in a periplasmic loop that folds inwards into the β-barrel. It is possible that this constricts the channel, as is observed in many outer membrane proteins.^2324^ Nevertheless, additional research on the matter is required to assess the function of Omp25 and the potential effect of NbH7.^17810^ In contrast, Omp25 appears to remain exposed and conserved across diverse isolates, which makes it a particularly interesting therapeutic or diagnostic target.

The conservation of Omp25 within *A. baumannii*, combined with the specificity of NbH7, supports its potential for diagnostic and therapeutic applications. This potential was explored through immunomagnetic capture of *A. baumannii* in mixed bacterial samples. Although the minimum bacterial concentration in the capturing assays leading to cluster formation was significantly lower than in other studies (10^2^ cells/mL^15^ or 10^4^ cells/mL^16^), the assay showed good discrimination between samples with and without *A. baumannii* bacteria. These results provide a proof-of-principle for cell enrichment, even for challenging multi-species samples including *E. coli*. Due to the presence of *E. coli* in the human gut microbiome^25^ and its link to antimicrobial resistance^26^, the lack of interaction between NbH7 and *E. coli* confirms its applicability as a precision-targeting tool.

The use of NbH7 as a diagnostic tool is of high potential. In this study, we demonstrate proof of concept for NbH7 as a method for targeted detection and bacterial enrichment of *A. baumannii* within a complex microbial environment. Although further optimization is needed, this feature of the Nb could enhance diagnostic accuracy and speed, in line with efforts to achieve early diagnosis capabilities. The binding specificity of NbH7 could be used to develop rapid diagnostic tests, possibly as biosensors or lateral flow assays, enabling quick identification of multidrug-resistant bacterial infections and supporting timely clinical decisions. Another potential application of NbH7 is to use the Nb to direct so-called ‘warheads’ to the bacteria in a specific way. For instance, these warheads could be nanoparticles infused with antimicrobial compounds. By conjugating NbH7 to antimicrobial agents, with all the advantages of Nbs, it would be possible to deliver them directly to bacteria, thereby increasing local drug concentration at the infection site and enhancing their efficacy through increased on-target retention time. Such targeted therapy could reduce the overall dosage required, minimize side effects, and decrease the selective pressure for the development of further resistance. Concerning diagnostics, assuming NbH7 is merely a binder, mutation frequency would not be elevated due to the lack of pressure on the bacteria. In addition, concerning targeted therapies, the presence of an identical Omp25 across the diverse tested *A. baumannii* and two other pathogenic species might indicate a need for the bacterium to conserve this protein and its unidentified function, but further testing is warranted. Although the tested isolate panel captures diversity in capsule type, it does not cover the impressive diversity of capsule types identified in *A. baumannii*.

In summary, we identify the outer membrane protein Omp25 as a conserved and accessible surface target in the highly diverse pathogen *A. baumannii* and demonstrate that the nanobody NbH7 selectively binds it across diverse *A. baumannii* isolates and two closely related species. The structural characterization of the Omp25-NbH7 complex provides molecular insights into their interaction, while successful immunomagnetic capture demonstrates the diagnostic potential of NbH7 in mixed samples.

Our findings highlight the potential of Omp25 and NbH7 as a starting point for the development of fast diagnostic tools and targeted therapeutic strategies against *A. baumannii*. Such Nb- or Omp25-based strategies could significantly impact the management of multidrug-resistant infections, addressing a critical need in contemporary healthcare. More broadly, this approach offers a versatile framework for developing Nb-based therapeutic or diagnostic strategies against other bacterial pathogens.

## MATERIAL & METHODS

### Bacterial strains and growth conditions

A collection of three classically used (AB5075-VUB - CP070362; AB5075-VUB-*itrA::*IS*Aba13* - CP070358; ATCC17978-VUB), three clinically isolated *A. baumannii* strains (AB3-VUB; AB180-VUB; AB220-VUB), three *Acinetobacter* spp. isolated from environmental sources (*A. calcoaceticus, A. junii* and *A. pittii*), *E. coli* strains (*E. coli* S17, ATCC 25922) and one *K. pneumoniae* strain were used in this study. The bacterial cultures were grown as described in Breine et al. (2023).^12^ In short, a single clone was used to start the cultures, which were grown for 17 h at 37°C under agitation (165 rpm) in low salt broth (Luria-Bertani formulation, Duchefa Biochemie). The bacteria were then 1:100 diluted in fresh LB medium and grown to an OD_600_ of 1 to obtain exponential phase bacteria. Both the stationary phase bacteria (from the overnight culture) and the exponential phase bacteria were collected by centrifugation (8000 g) and resuspended in phosphate-buffered saline (PBS) to obtain an OD_600_ of 3 (10^9^ CFU/ml) for further use.

### Nanobody discovery, purification and labelling

The technical aspects of the generation of the nanobody library are described in Breine et al. (2023).^12^ The nanobodies are expressed in and purified from the periplasm of *E. coli* as described elsewhere.^27^ In short, the coding sequence of the nanobody is cloned into the phagemid vector pMESy4, in frame with a N-terminal periplasmic leader sequence and a C-terminal 6-His-EPEA tag. This plasmid is then transformed into *E. coli* WK6 cells and the cells are grown in Terrific Broth (Duchefa Biochemie) and periplasmic expression of the nanobodies is induced by IPTG overnight. The cells are harvested by centrifugation (10 min, 6000g) and resuspended in a periplasmic extraction buffer (Tris-HCl pH 8, 20% sucrose, 500 mM NaCl, 2 mM EDTA, 0.1 mg/ml lysozyme, 50 μg/ml DNase, protease inhibitors) for 1h. The suspension is centrifuged (45 min, 17700 g) to obtain the periplasmic extract. The nanobodies are purified from the periplasmic extract by Immobilized Metal Affinity Chromatography. For further tests, the nanobodies were dialyzed to PBS overnight.

To fluorescently label the nanobodies, they were incubated with a 3-fold molar excess of DyLight 650 nm NHS ester (ex:652/em:672, ThermoFisher) for 1 h, after which the reaction was quenched with 50 mM Tris buffer, followed by overnight dialysis in PBS.

### Fluorescence microscopy

The binding of the Nbs was tested by incubating equal volumes of the bacteria (a 1:1 mix of exponential and stationary phase bacteria) at approximately 10^5^ CFU with 10 μM Nbs. The unbound Nbs were removed after 30 min incubation under agitation (165 rpm) by 3 centrifugation steps (2 min, 8000 g). The bacteria were spotted on a 1.5% agarose pad (Thermo Scientific Gene Frame) for visualization with the microscope. Images were acquired using a Leica DMi8 fluorescence microscope with a DFC7000 GT camera (Leica Microsystems CMS GmbH). For one assay (**Figure S2**), an AB5075-VUB strain expressing cytoplasmic GFP was used as well. The fluorescent images were acquired with a Leica FRAP450 and Y5 filter set. The raw data was processed by using ImageJ software where brightness was adjusted equally for all fluorescence micrographs within one experiment. All data was collected in biological triplicate.

### Pull-down

For the pull-down experiment, approximately 10^10^ CFU of AB5075-VUB and AB5075-VUB-*itrA*::IS*Aba13* were lysed by PBS supplemented with 1% n-Dodecyl-β-D-maltoside, 300 mM NaCl, 10 mM imidazole, 50 μg/ml DNase and 0.1 mg/ml lysozyme for 2 h. The lysed cultures were incubated with 200 μM Nb, including a control Nb (CtrlNb), for 1 h at 37°C under agitation (165 rpm). This cell lysate mixture was loaded over His SpinTrap columns (Cytiva) according to the manufacturer’s instructions. The Nb-target complex could be pulled out of this mixture due to the C terminal hexahistidin tag of the Nb. The columns were washed with PBS supplemented with 0.03% DDM, 20 mM imidazole and 300 mM NaCl, and eluted with PBS supplemented with 0.03% DDM, 500 mM imidazole and 300 mM NaCl. The eluted samples were consequently analyzed by sodium dodecyl sulfate–polyacrylamide gel electrophoresis (SDS-PAGE). The band visible around 25 kDa in the NbH7 sample, but not the CtrlNb samples, was excised. The VIB Proteomics Core facility identified the proteins present in this band by liquid chromatography-tandem mass spectrometry (LC-MS/MS).^28^ All samples were searched separately using the MaxQuant algorithm (version 2.0.1.0) with mainly default search settings, including a false discovery rate set at 1% on PSM, peptide and protein level. Spectra were searched against the *Acinetobacter baumannii* taxid470 UniProt database^29^, a control protein sequence and the NbH7 protein sequence. The data were ordered according to Intensity-Based Absolute Quantification, which revealed the most abundant proteins present in the sample.

### Construction of AB5075C+Δ*omp25* and AB5075C-Δ*omp25* strains

The AB5075C*+*Δ*omp25* and AB5075C-Δ*omp25* strains were generated by making a construct of the *sacB-aaC* selection/counterselection cassette flanked by 2 kb up and downstream homologous regions of the *omp25* gene of AB5075-VUB, followed by transformation to both bacterial strains as described in Whiteway *et al*.^14^

### Recombinant expression of AB5075-VUB Omp25 in the OM of *E. coli*

For recombinant expression of AB5075-VUB Omp25 in the outer membrane (OM) of *E. coli*, the signal peptide (SP) coding sequence of *E. coli* OmpA was used. By means of PCR, the open reading frame of AB5075-VUB Omp25, excluding its endogenous SP coding sequence, was cloned into a pASK-IBA43 plus vector downstream from a N-terminal streptavidin tag, a short glycine-serine linker, and a TEV cleavage site to construct pASK-Omp25. The pASK-Omp25 plasmid was transformed into *E. coli* BL21 (DE3) pLysS Δ*tonA* cells. An overnight culture started from a transformant was used to inoculate LB supplemented with ampicillin and 2mM MgCl2 and 0,1% (w/v) glucose. The culture was grown to an OD_600_ of 0.5-0.7, after which Omp25 expression was induced with 0.2 μg/ml anhydrotetracycline hydrochloride. After 3h, the cells were either tested for binding of NbH7 with fluorescence microscopy or used for purification of Omp25.

### Omp25 protein purification

The *E. coli* BL21 (DE3) pLysS Δ*tonA* cells were harvested by centrifugation (10 min, 6000 g) and resuspended in lysis buffer containing 50 mM Tris-HCl pH 8.0, 300 mM NaCl, 10% glycerol, 1 mM β-mercaptoethanol, DNase and an antiprotease cocktail. The cell suspension was run through a LM10 Microfluidizer (Microfluidics™) at 15000 psi to lyse the cells. The cell lysate was centrifuged (45 min, 48000 g) to obtain the membrane fraction. n-Octylpolyoxyethylene (n-OPOE) was used to solubilize Omp25. The pellet was resuspended in buffer containing 50 mM Tris-HCl pH 8.0, 300 mM NaCl, 3% OPOE and an antiprotease cocktail. After 1h incubation at 4°C the membrane extract was centrifuged (45 min, 48000 g) to obtain the solubilized membrane fraction containing the strep-tagged protein. This membrane fraction was loaded onto a Strep-Tactin®XT 4Flow® cartridges (IBA) and washed with 100 mM Tris-HCl pH 8.0, 150 mM NaCl, 1% OPOE and 1 mM EDTA. Protein that was used for BLI analysis was eluted with 100 mM Tris-HCl pH 8.0, 150 mM NaCl, 1% OPOE, 1 mM EDTA, 50 mM biotin. The eluted protein was analysed by SDS-PAGE to assess the purity, then concentrated by using an AMICON 10 kDa MWCO and flash frozen in liquid nitrogen to be stored at -80°C. Protein that was used for crystallization was first detergent-exchanged on column by washing with 100 mM Tris-HCl pH=8, 150 mM NaCl, 5 mM LDAO, 1 mM EDTA and then eluted by 100 mM Tris-HCl pH=8, 150 mM NaCl, 5 mM LDAO, 1 mM EDTA, 50 mM biotin. Eluted protein was loaded onto a Superdex 100 16/60 size-exclusion column (GE life Sciences) that was equilibrated with SEC buffer containing 100 mM Tris-HCl pH=8, 150 mM NaCl, 2 mM LDAO. The fractions corresponding to Omp25 were pooled and consequently used for crystallization.

### Bio-layer interferometry (BLI)

To determine the affinity of NbH7 and its target Omp25, biolayer interferometry (BLI) was done on an Octet® R8 instrument (Sartorius). All measurements were done at 30°C in 50 mM Tris (pH 8), 150 mM NaCl, 1% n-OPOE and 0.1% BSA. The nanobodies were loaded on Ni-NTA sensors (Sartorius, #18-5101) at a concentration of 10-80 μg/ml for 120 s to obtain approximately 1 nm signal increase. After a baseline of 480 s, the association was measured for 100 s by dipping the sensors in a two-fold dilution series starting at 80 μM Omp25. The dissociation was measured for 600 s. To monitor aspecific binding, a control Nb (CtrlNb) was tested in parallel in each measurement against the highest concentration of Omp25 (80 μM) **(Figure S9)**. This aspecific binding signal was subtracted from each binding signal before further calculations. The R_eq_-values were plotted against the Omp25 concentration and fitted on a one-site specific binding curve using Graphpad Prism to determine the equilibrium dissociation constant (K_D_ ± standard error). The binding curves of the lowest Omp25 concentrations, 2.5, 5 and 10 nM, could not be fitted and calculated. The R_eq_-values were calculated using Octet® Analysis Studio software (version 13.0.2.46) and the graphs were plotted using Graphpad Prism (version 9.0.1).

### Crystallization, structure determination and analysis

The crystallization screens were set up with a mix of freshly purified Omp25 with 1.2-fold molar excess of NbH7. The mixture was concentrated to 35 mg/ml by using an AMICON 3 kDa MWCO. Crystals were obtained by using the sitting drop vapor diffusion method and a Mosquito nanoliter-dispensing robot at room temperature (TTP Labtech, Melbourn, UK). After 3 months, hexagonal-shaped crystals appeared after in a crystallization buffer containing 0.2 M calcium chloride dihydrate, 0.1 M Tris-HCl pH 8.0 and 44 % v/v PEG 400. After supplementing the buffer with 10% glycerol, the crystals were mounted in nylon loops and flash-cooled in liquid nitrogen. The X-ray diffraction data was collected at 100 K using the Beamline Proxima 2A (wavelength = 0.9801 Å) at the Soleil Synchrotron (Gif-sur-Yvette, France). The diffraction data was processed using autoPROC^30^ software at 2.91 Å in R3 with unit-cell dimensions of a = 190.55, b = 190.55, c = 231.103, ⍰ = 90.0, β = 90.0, γ = 120.0. To determine the crystal structure, molecular replacement was done using phaser from the phenix suite^31^ with as search models the NbH7 and Omp25 structures predicted by AlphaFold2^32^. Refinement of the structure was done by iterative cycles of manual model building with COOT^33^ and refinement with phenix.refine^31^ until R values of R_work_/R_free_ of 0.24/0.27 were obtained. More crystallographic parameters can be found in supplementary table 2 (**Table S3**). Figures of the structures were generated with ChimeraX^34^ and the NbH7-Omp25 interaction was determined by ChimeraX^34^ and PDBePISA^35^

### Genomic analysis *omp25* conservation

In January 2026, the NCBI’s RefSeq ^36^ was assessed for complete genomes of *A. baumannii* (n=1035) in order to avoid a bias from the fragmented draft-genome assemblies. The genomes were screened for the presence of *omp25* of AB5075-VUB (CP070362, locus JWB56_17755). The nucleotide identity of the putative *omp31* gene was determined by screening the genomes obtained from the NCBI’s RefSeq for the gene encoding the putative *omp31* of *A. baumannii* ZW85-1 (CP006768, locus P795_0840. Cutoff values for nucleotide identity and coverage were both 80%.

### Immunomagnetic capture using NbH7-coated magnetic particles (MP) and electrical analysis of MP-*A. baumannii* clusters

The immobilization of NbH7 onto the surface of the MP was achieved using the His-tag present in the nanobody, following the Invitrogen protocol. In short, the Invitrogen 1 μm diameter MPs Dynabeads™ His-tag beads were resuspended in a binding/washing buffer (50 mM Sodium Phosphate, pH 8.0, 300 mM NaCl, 0.01% Tween 20). The suspension with 2 μg/mL NbH7 nanobody was incubated overnight on a roller. Finally, the MPs were washed 5 times with the binding/washing buffer and resuspended in measurement buffer to a concentration of 6 mg beads/mL.

A suspension of 6 μL NbH7-coated MPs from the stock solution was resuspended in 1.5 mL PBS (20 mM, pH 7.4, 0.02% Tween 20, 0.1 mg/mL BSA) inside the measurement chamber^37^. The MPs were magnetically collected in the vicinity of the measurement electrode. Magnetic oscillations of the MPs cloud are induced to analyse the baseline signal of the MPs alone. Concentrations of *A. baumannii* 10^5^, 5 10^5^ and 10^6^ cells/mL, as well as 10^6^ cells/mL of non-target *E. coli*, individually or as mixtures, added to the chamber, were affinity bound to the MPs redispersed in solution. After 15 minutes of incubation, the MPs (unbound and the MP-bacteria clusters) were magnetically collected on the measurement electrode. The electrical response was monitored while clusters containing MP cloud underwent controlled magnetic oscillations. The amplitudes of impedance oscillations, resulting from the magnetophoretic movement of the MP cloud without or with various cell concentrations are measured at two chosen AC frequencies (*Lf* =455 kHz and Hf=5.5 MHz, respectively).^16^

For each cell concentration, the high and low AC frequency datasets are processed by calculating the quadratic mean,

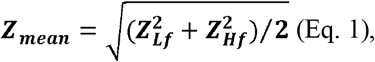

where Z_Lf_ and Z_Hf_ represent the normalized electrical impedances, at the low and high AC frequencies, respectively. These data, are normalized against control, i.e. *Z*_*meanMP*_ (measured on MPs per se, without bacteria), using the formula:

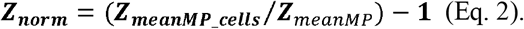

The normalized values (Z_***norm***_) according to Eq. 2 are expressed as mean ± standard deviation from three independent replicates for each concentration of target bacteria (*A. baumannii*) as monoculture or mixed with nontarget *E. coli*.

## Acknowledgements

A.B., A.V., C.V.d.H., and H.R. thank the Research Foundation−Flanders (FWO Vlaanderen) for providing a PhD SB fellowship to A.B. (File number 77258) and a junior postdoctoral fellowship to A.V. (File number 1287223N). C.V.d.H. acknowledges the Vlaams Instituut voor Biotechnologie (VIB) for their support. E.P. and J.S. acknowledge the support and the use of resources of Instruct-ERIC, part of the European Strategy Forum on Research Infrastructures (ESFRI), and the Research Foundation− Flanders (FWO) for their support to the Nanobody discovery and Nele Buys for the technical assistance during Nanobody discovery. M.G., D.T., S.D. and E.G. thank the Romanian Ministry for Education and Research, grant PNRR-III-C9-2023-I8, contract CF129-31.07.2023. This work was also supported by the European BAXERNA 2.0 project (101080544).

## Contributions

C.V.d.H., E.P., J.S., and H.R. generated the nanobody library. A.B. and C.V.d.H. identified the binding nanobody. A.B. performed the microbiological work, microscopy experiments, produced the knockout mutants, produced the recombinant expression strains and purified and co-crystallized the proteins, performed the biochemical study of the complex, and analyzed the data. S.V.M. performed the expression and purification of the nanobodies as well as the recombinantly produced protein. A.B., E.J. and H.R. performed the structural study of the complex. A.V. performed the genomic analysis. C.V.d.H. and A.B. conceptualized the study. C.V.d.H. supervised the study. A.B., E.J. and C.V.d.H. wrote the paper, with contributions from all authors. M.G. conceptualization, editing, and reviewing, D.T. microbiological work, S.D. microscopy experiments, magnetic particles functionalization, impedance experiments, data processing, E.G. conceptualization, data analysis reviewing.

## Ethics declarations / competing interests

C.V.d.H. and A.B. are named as inventors on a patent describing the use of NbH7 to target *Acinetobacter baumannii*.

## Supplementary information

See additional document.

