## Supplementary material for "A novel nanobody-based approach for targeting heterogeneous *Acinetobacter baumannii* isolates and closely related pathogenic *Acinetobacter* spp": Breine et al Supplementary information.docx

**Figure S1**

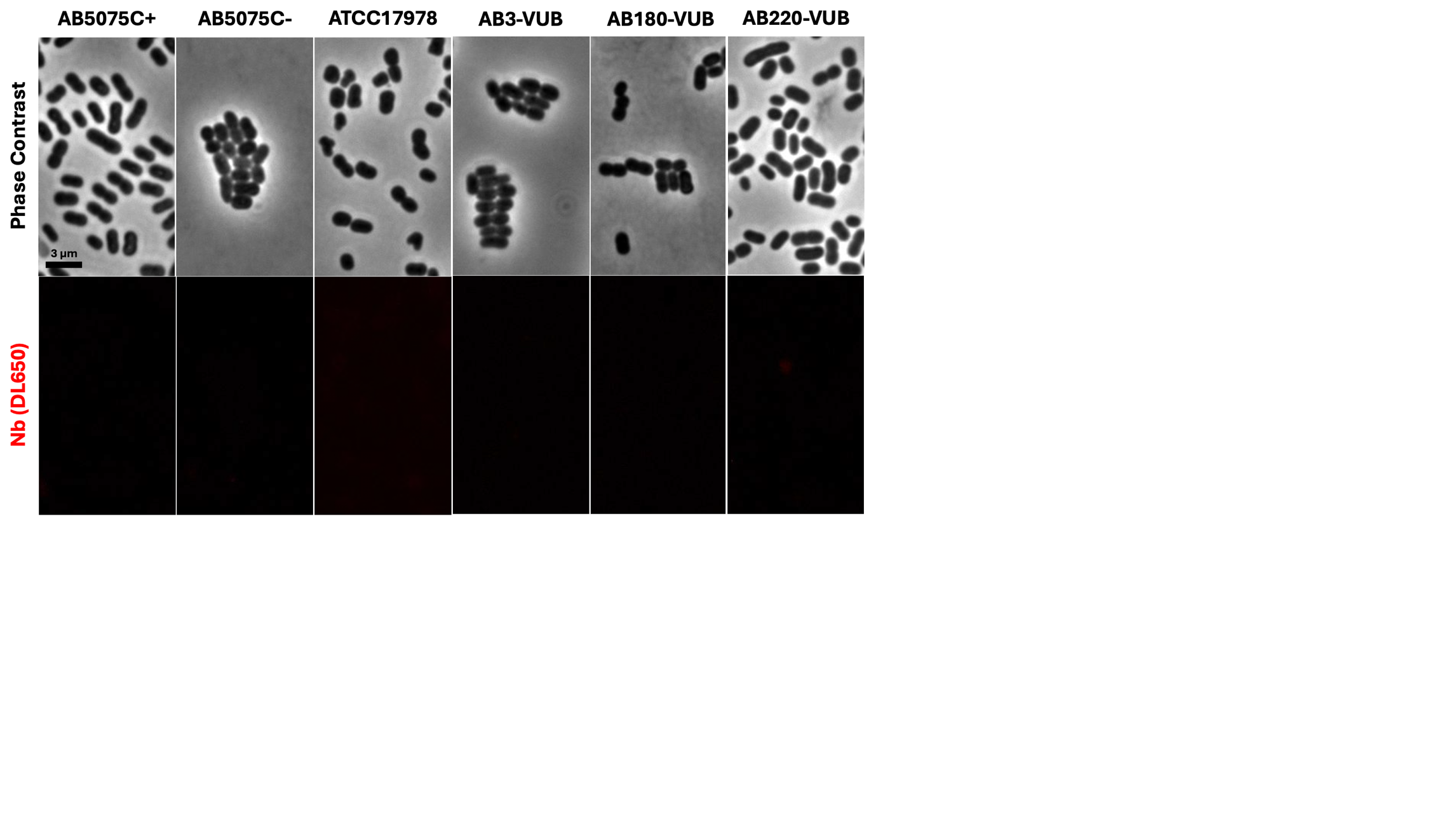
**Figure S1.** **The control Nb (CtrlNb) does not show any binding to *A. baumannii*.** In each experiment, a control Nb that does not bind *A. baumannii*, was also labelled with DL650 and tested in the exact same conditions by fluorescence microscopy. Images were taken and analyzed in the same manner as for the other Nbs. This is an overview of those micrographs on the tested *A. baumannii* strains. No binding was observed. All data was collected in biological triplicate.

**Figure S2**

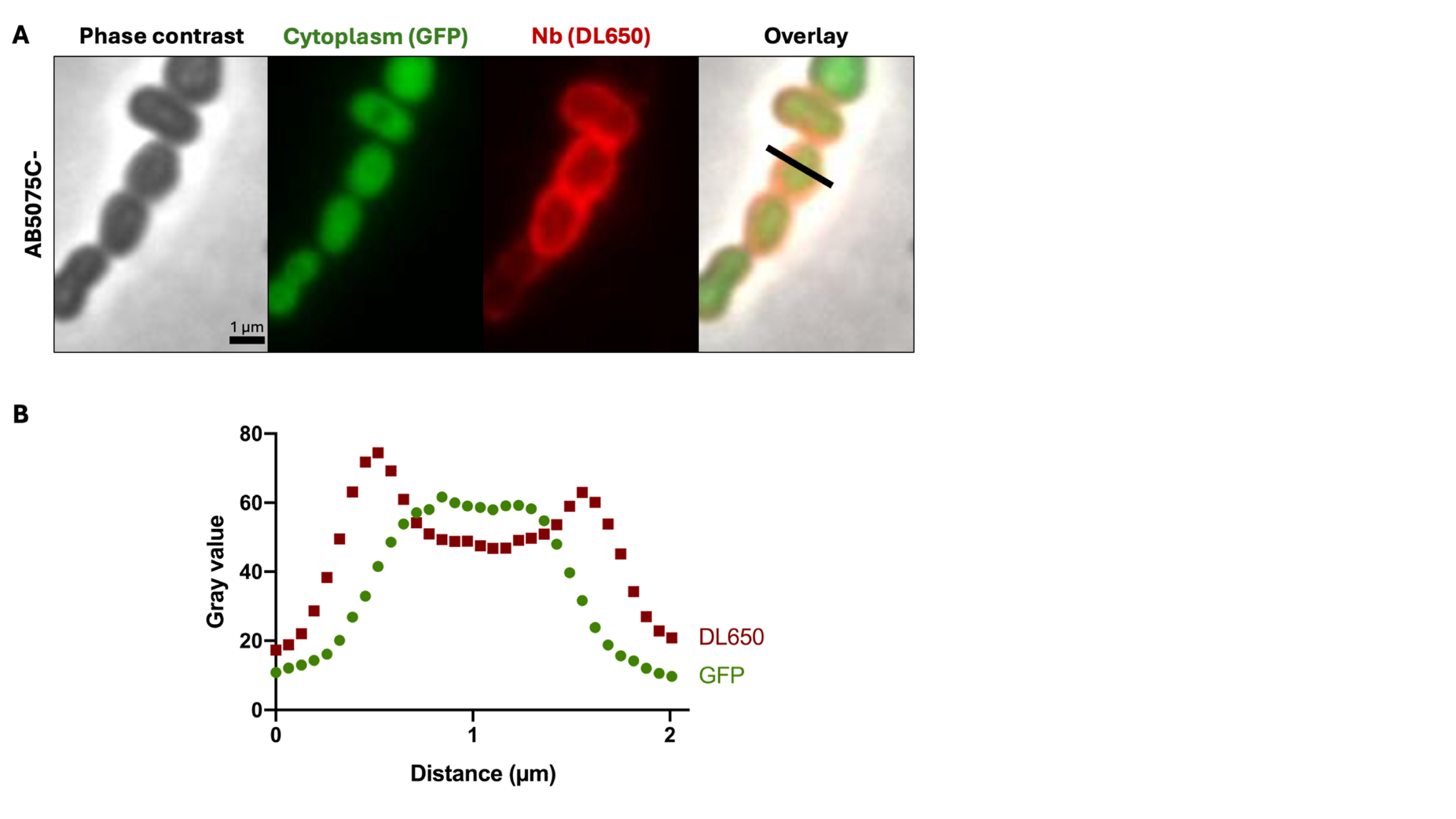

**Figure S2. NbH7 binds the membrane of AB5075C-. (A)** Fluorescence micrograph of AB5075C- cells expressing GFP in the cytoplasm. The bacteria are bound by NbH7 that was fluorescently labelled with DyLight 650 nm (DL650). An intensity profile was measured across a representative bacterial cell (black line in the overlay image) and is plotted in (**B).** The intensity profiles of GFP and DL650 show a cytoplasmic labelling profile for GFP and a membrane labelling profile for NbH7 (DL650). Graph generated by ImageJ and Graphpad Prism. All data were collected in biological triplicate.

**Figure S3**

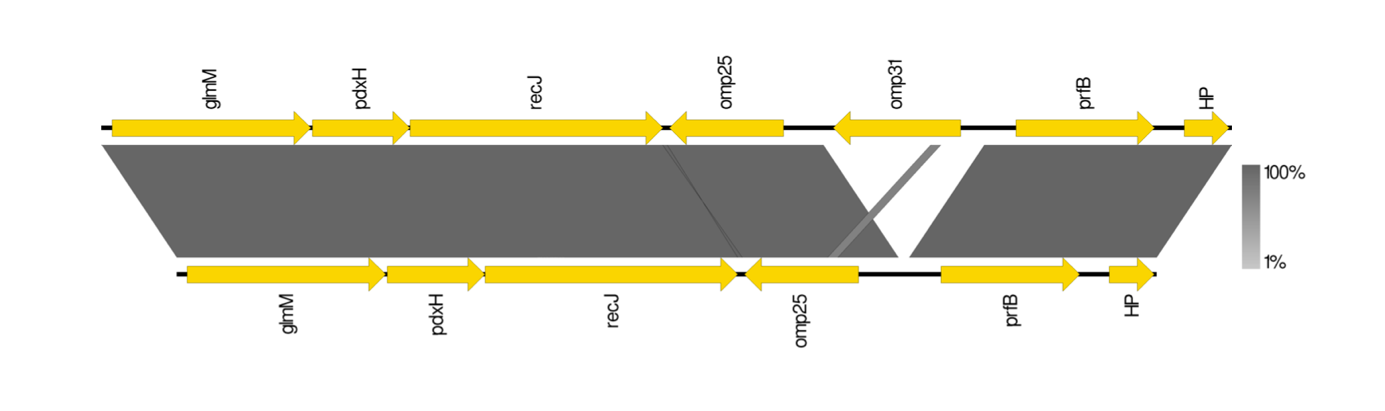

**Figure S3 - The analysis of the genetic environment of the *omp25* revealed two variants of organization.** The genetic environment of the *omp25* is conserved, however in only 121 out of 1035 complete genomes of *A. baumannii* (11.7%), a gene encoding a putative outer membrane protein of 31kDa (designated *omp31*) was detected upstream of the *omp25* gene.

**Figure S4**

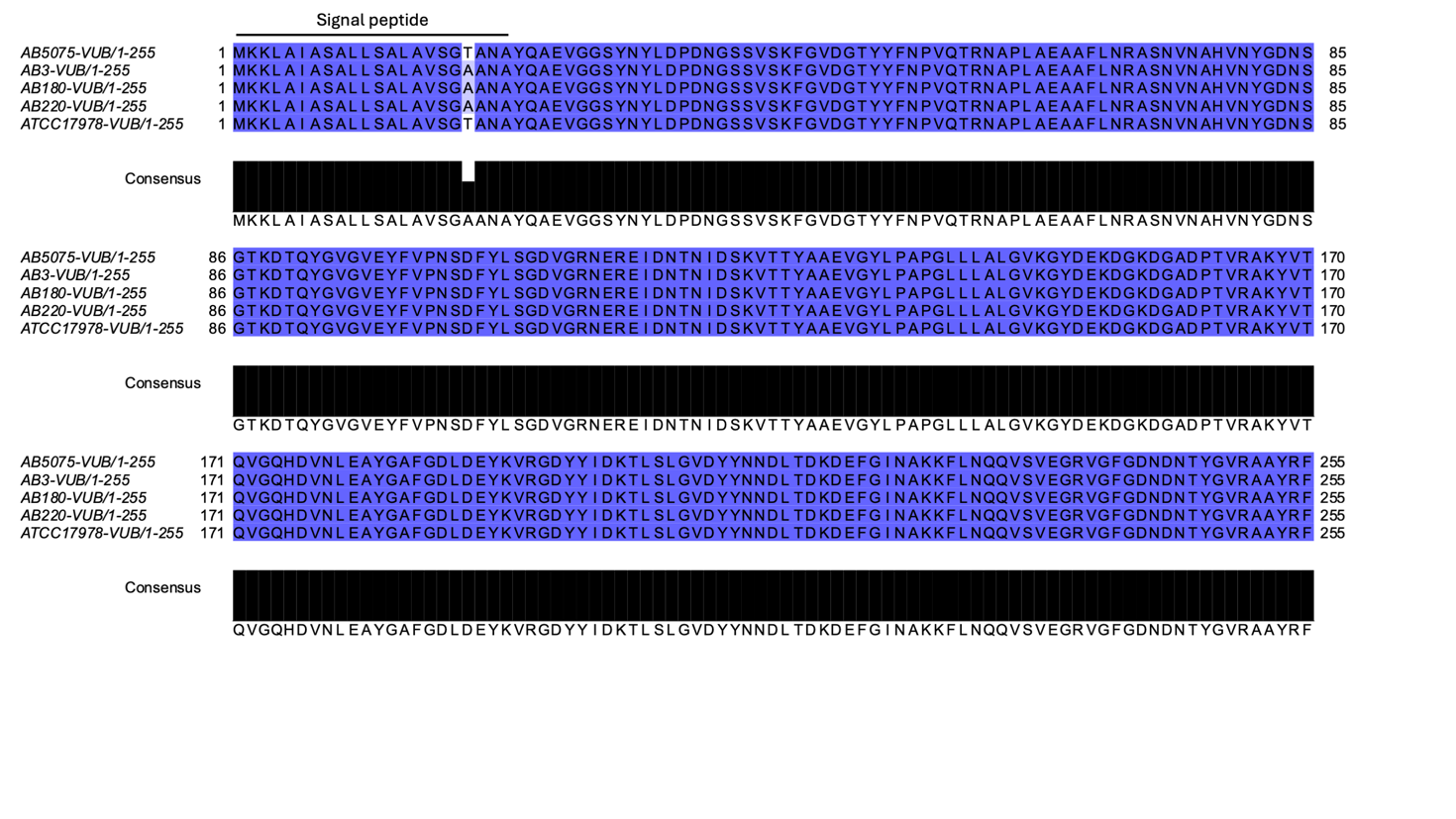

**Figure S4. Multiple sequence alignment of AB5075-VUB Omp25 to all the tested *A. baumannii* strains.** Excluding the signal peptide, which is not present in the final protein, the Omp25 protein sequences are identical. The alignment was generated using Clustal Omega and Jalview software.

**Tabel S2**

|  | **% ID with AB5075 Omp25** | **Accession number** | **Length (#aa)** |
| --- | --- | --- | --- |
| ***Acinetobacter pittii*** | 94,14 | WP_337077925.1 | 255 |
|  | 96,57 | RQL72781.1 | 255 |
|  | 95,71 | WP_063097693.1 | 255 |
|  | 94,85 | WP_216946930.1 | 255 |
| ***Acinetobacter calcoaceticus*** | 96,14 | WP_016139874.1 | 255 |
|  | 95,71 | WP_004639960.1 | 255 |
|  | 95,28 | WP_199966399.1 | 255 |
|  | 94,85 | VAX43036.1 | 255 |
|  | 93,13 | CAI3136485.1 | 255 |
| ***Acinetobacter junii*** | 81,97 | WP_004951331.1 | 255 |
|  | 81,55 | RXS99878.1 | 255 |
|  | 81,12 | WP_262579446.1 | 255 |

**Table S2. Homologous Omp25 sequences in *A. pittii*, *A. calcoaceticus* and *A. junii*.** A Protein BLAST was done using AB5075-VUB Omp25 against a non-redundant protein sequences database in *Acinetobacter* (taxid: 469), excluding *Acinetobacter baumannii* (taxid: 470). The results were filtered for each species separately and one representative of differing sequence identity (% ID) was selected for each hit with the same amino acid sequence length (#aa): 255. For each hit, the accession number (GenBank) is given.

**Figure S5**
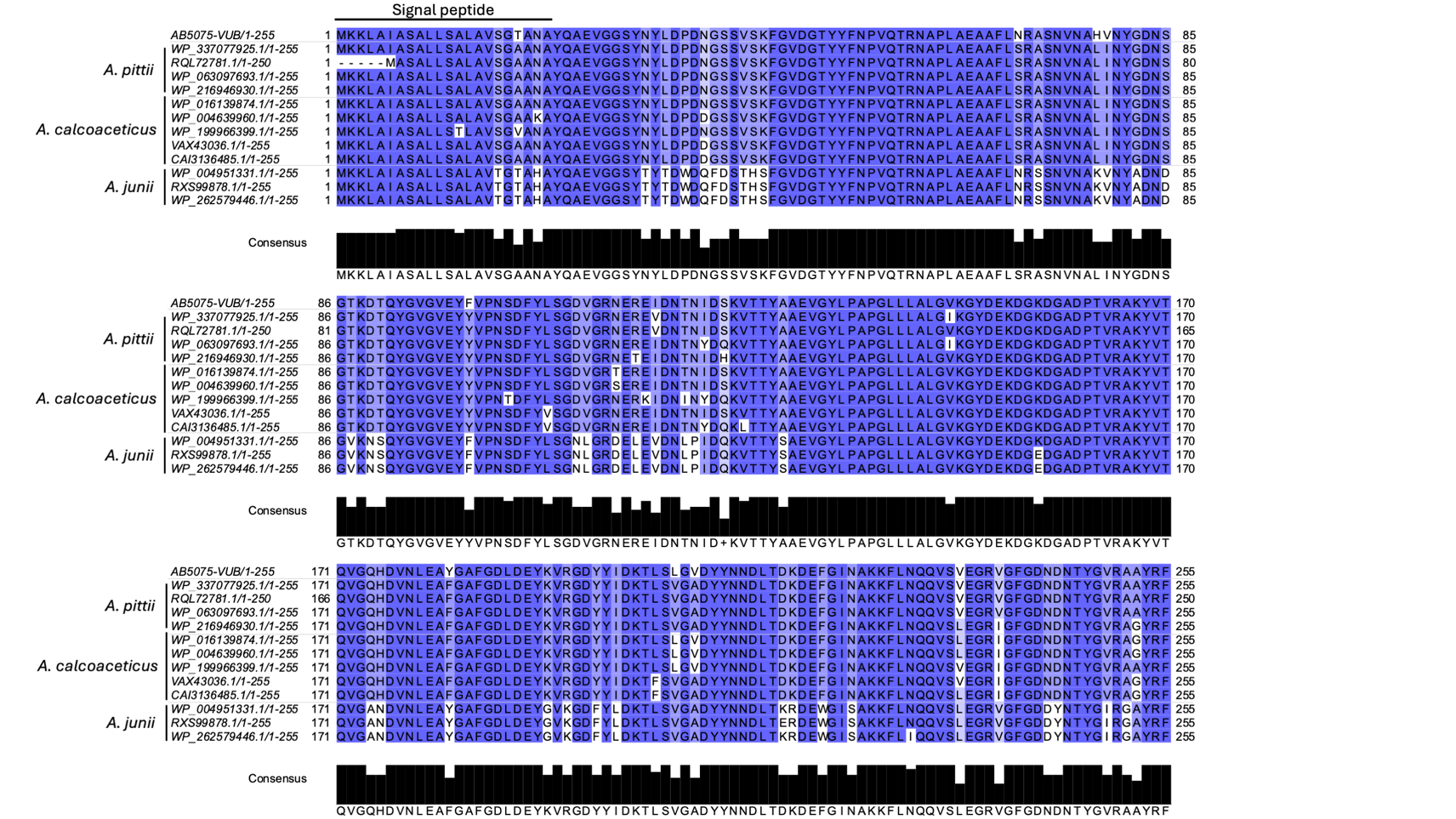

**Figure S5. Multiple sequence alignment of selected *A. pittii*, *A. calcoaceticus* and *A. junii* Omp25 amino acid sequences against AB5075-VUB Omp25.** The less conserved amino acids are lighter or not colored, which is also represented in the height of the bars in the consensus graph. More information on the strains is provided in **Table S2**. The alignment was generated using ClustalWS and Jalview software.

**Figure S6**

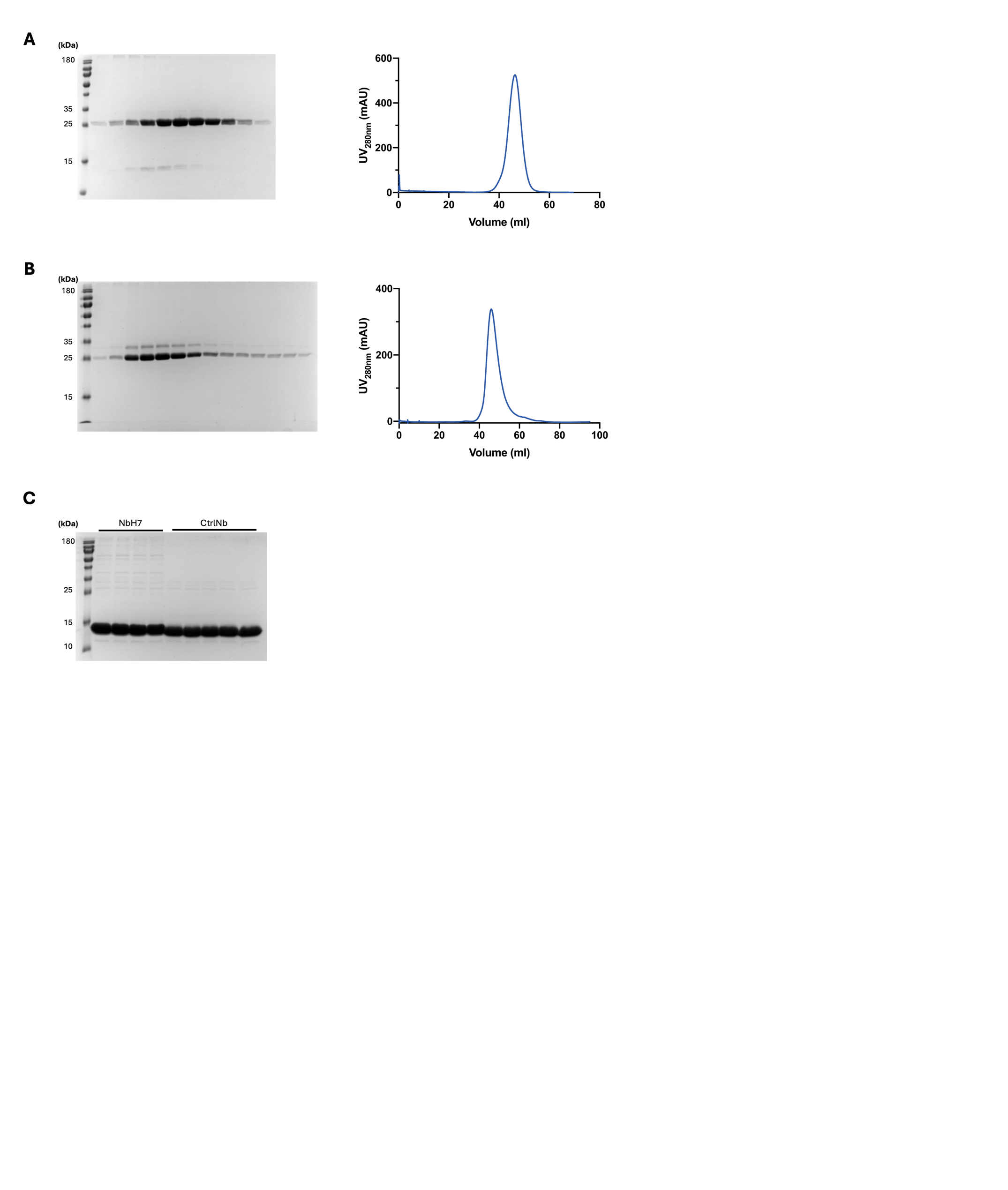

**B**

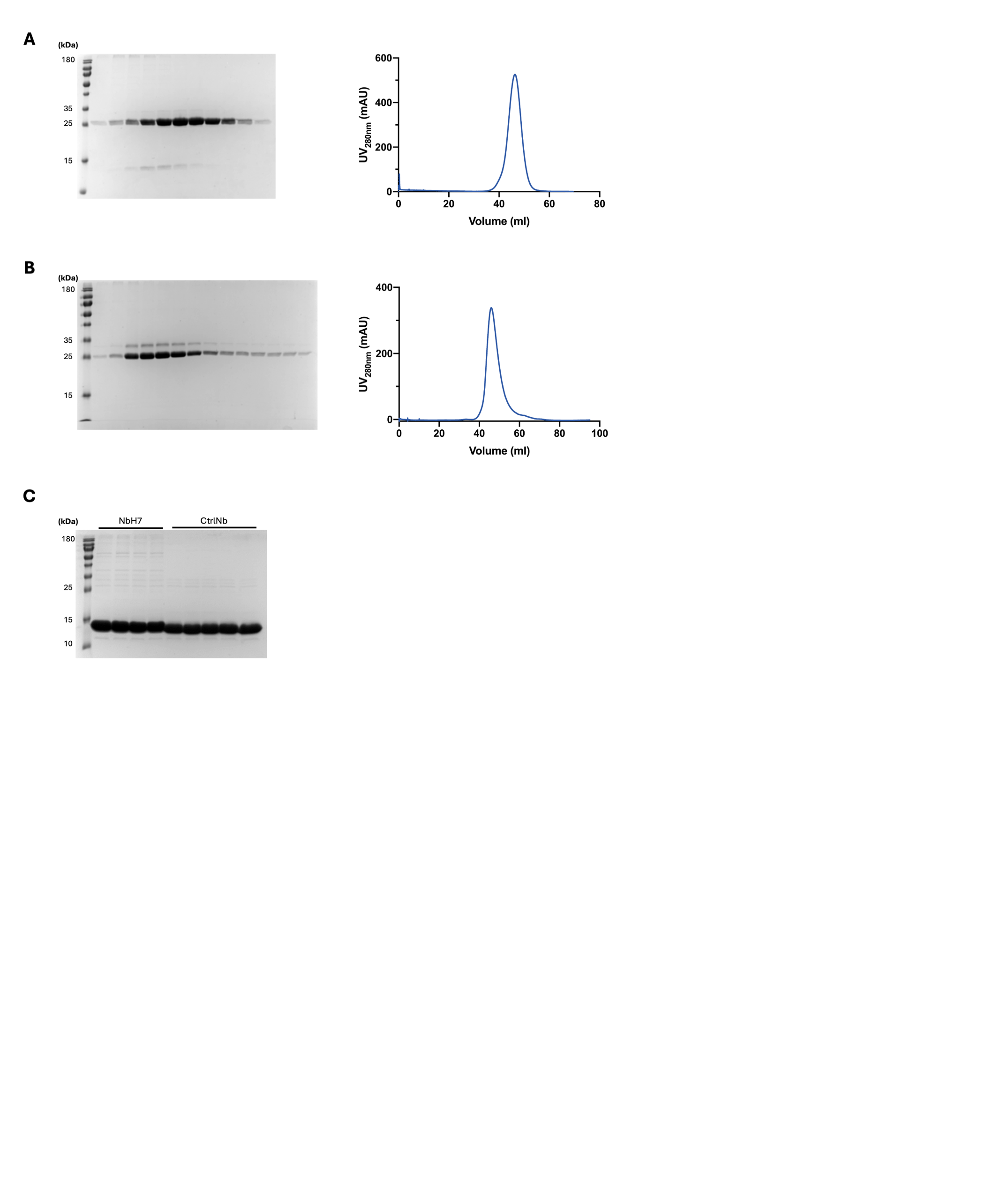

**Figure S6. Quality analysis of the purified proteins. (A)** SDS-PAGE and Size Exclusion Chromatography (SEC) profile of purified Omp25 using OPOE**.** SDS-PAGE samples correspond to fractions of the size exclusion chromatogram. (**B)** SDS-PAGE of the purified nanobodies.

**Figure S7**

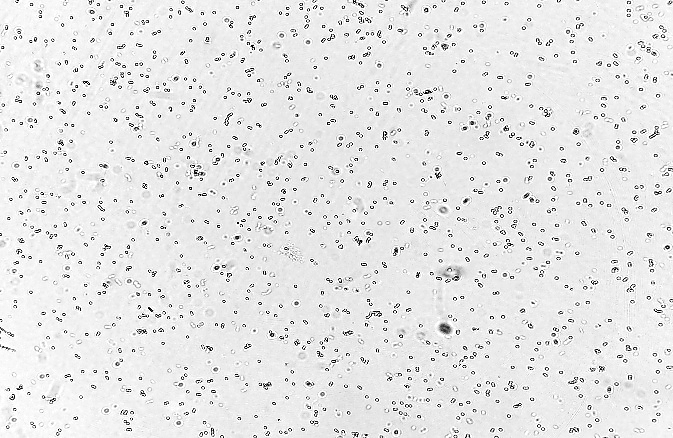

10 μm

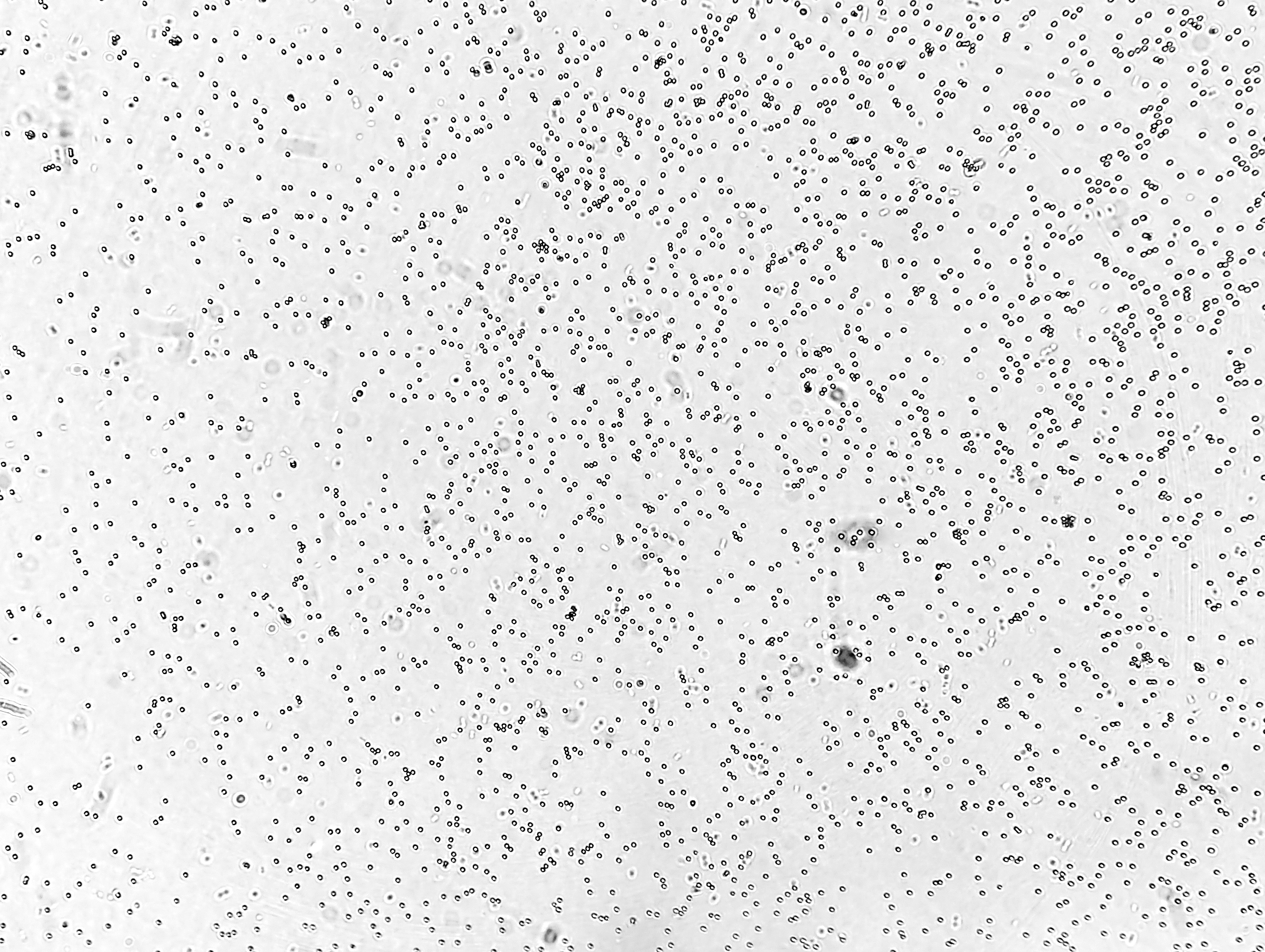

10 μm

A

B

**Figure S7. Non-specific *A. baumannii*- MP interactions.** Transmission images of (**A)** MP functionalized with non-specific anti-*E. coli* antibody, (**B)** MP subjected to the full surface immobilization protocol without protein addition, after incubation with *A. baumannii* (~10⁹ cells/mL). No MP aggregation is visible, confirming that the aggregation is strictly dependent on the presence of NbH7 on the MP surface.

**Figure S8**

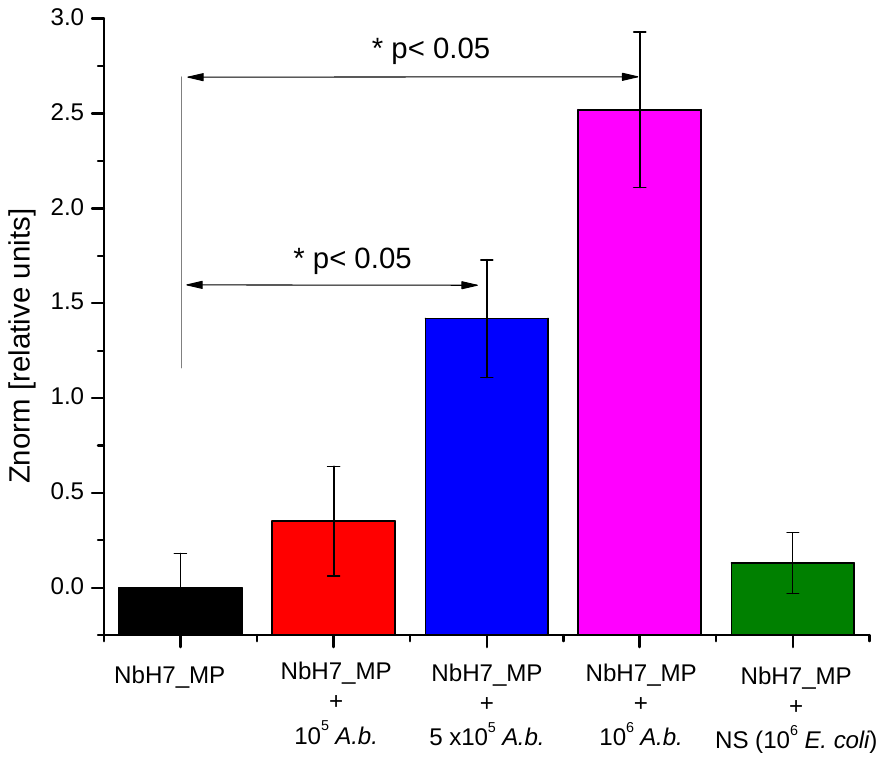
(**A**)

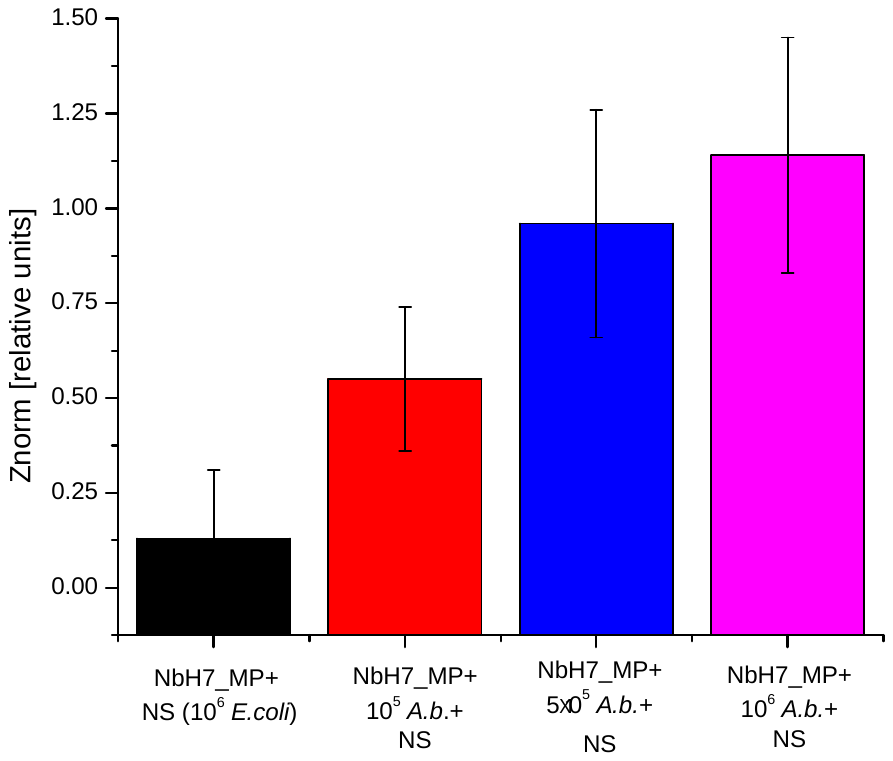
(B)

**Figure S8.** **Bar graphs of the normalized amplitudes of impedance oscillations indicative for specific aggregation of NbH7-coated MPs in the presence of *A. baumannii*.** (**A)** The concentration-dependent response for pure *A. Baumannii* suspensions. NS represents the response in the presence of 10^6^ cells/mL non-target bacteria (**B).** The concentration-dependent response for *A. Baumannii* suspensions in the presence of non-specific cells – NS 10^6^ cells/mL *E. coli* bacteria. Error bars correspond to the standard deviation for N=3 individual experiments.

**Figure S9**

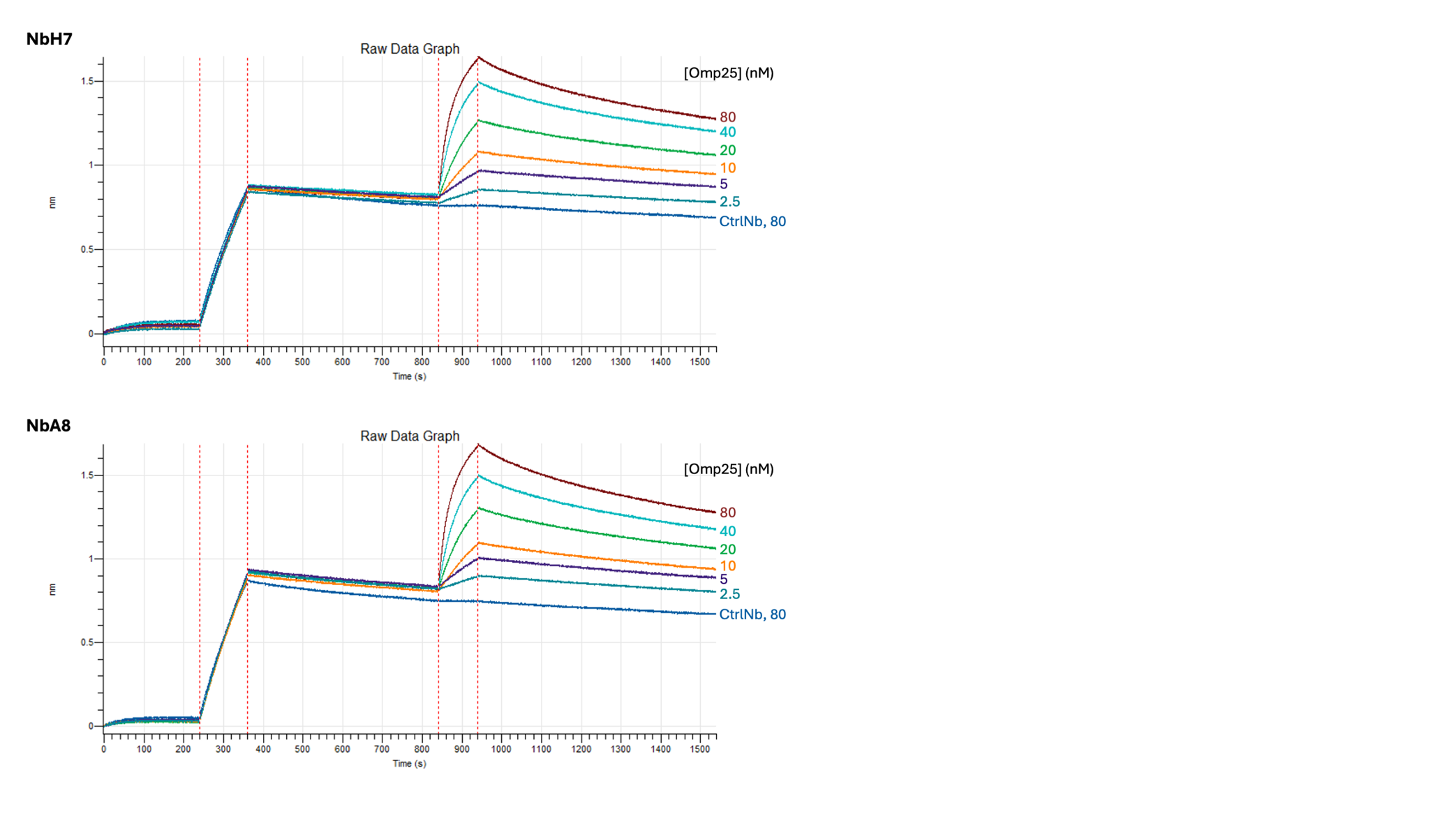

**Figure S9. Raw data of BLI measurements to determine the affinity of NbH7 for Omp25.** A control Nb (CtrlNb) was measured against the highest concentration of Omp25 to monitor aspecific binding. This nonspecific binding signal was subtracted from the binding signals of NbH7 before analysis of the measurement. All measurements were done at 30°C.

**Table S3. Crystallographic parameters of the Omp25-NbH7 dataset.** The values of the highest recorded resolution shell are given in parenthesis.

| **Data collection** | |
| --- | --- |
| Space group | R3 |
| Cell dimensions | |
| a, b, c (Å) | 190.55 190.55 231.103 |
| ⍺, β, γ (°) | 90.0 90.0 120.0 |
| Resolution (Å) | 20.01 - 2.87 (3.05 - 2.87) |
| Rmeas (%) | 38.1 (207.7) |
| I/σ (I) | 5.4 (1.4) |
| Completeness (%) | 94.7 (47.7) |
| Multiplicity | 9.0 (9.0) |
| Wilson B factor (Å^2^) | 51.90 |
| **Refinement** | |
| Resolution (Å) | 2.91 |
| Number of unique reflections | 62255 (3113) |
| R_work_/R_free_ | 0.24/0.27 |
| Number of protein atoms | 2814 |
| Average B-factor (Å^2^) | 59,72 |
| RMS deviations | |
| Bond length (Å) | 0.01 |
| Bond angles (°) | 1.79 |
| **PDB code** | Not known yet |

**Figure S10**

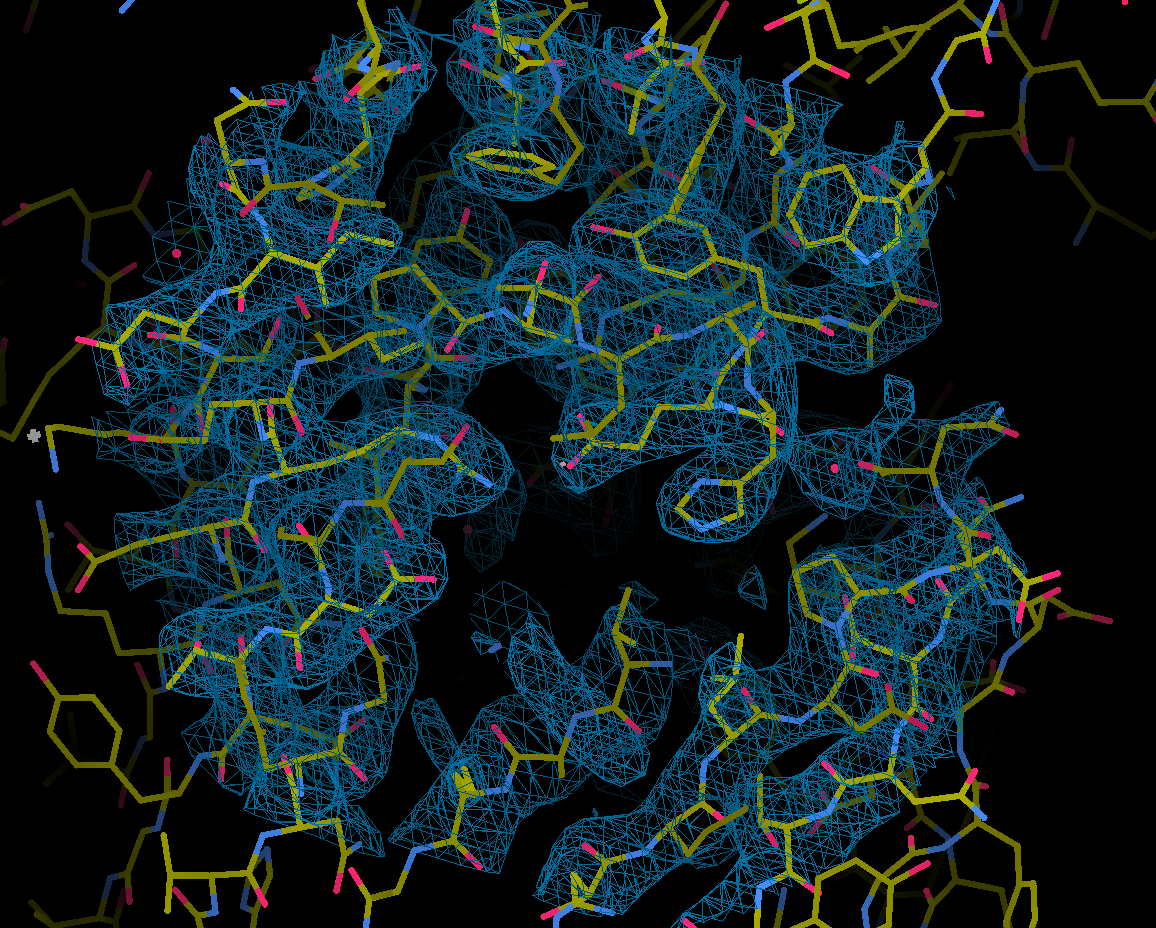

**Figure S10. Electron density map structure Omp25-NbH7.** Portion of the 2Fo-FC map at contour level of 1.2 σ of the interface between NbH7 and Omp25.
